# Biosphere functional integrity for people and Planet

**DOI:** 10.1101/2022.06.24.497294

**Authors:** Awaz Mohamed, Fabrice DeClerck, Peter H. Verburg, David Obura, Jesse F. Abrams, Noelia Zafra-Calvo, Juan Rocha, Natalia Estrada-Carmona, Alexander Fremier, Sarah K. Jones, Ina C. Meier, Ben Stewart-Koster

**Author notes:** Corresponding author: Awaz Mohamed.

## Abstract

Defining a safe and just biosphere space requires a synthetic scaleable measure of biosphere functional integrity to secure Nature’s Contributions to People (NCP). Using a systematic review of 153 peer-reviewed studies we estimated the minimum level of functional integrity needed to secure multiple critical NCP, including pollination, pest and disease control, water quality regulation, soil protection, recreation and natural hazards mitigation in human-modified landscapes. We characterise functional integrity by the quantity, quality and spatial configuration of (semi-)natural habitat within any landscape. We find that at least 20-25% of structurally complex and biologically diverse (semi-)natural habitat in each 1 km^2^ of land area is needed to maintain the supply of multiple NCP simultaneously. Exact quantity, quality and spatial configuration required is dependent on local context, and may differ for individual NCP. Today, about 50-60% of human-modified lands have less than 10% and 20% (semi-)natural habitat per 1 km^2^ respectively. These areas require immediate attention to regenerate functional integrity in order to secure ecological functioning in those landscapes.

---

Most attention in biodiversity conservation is given to halting the conversion of remaining natural ecosystems, the unique species they hold^1,2^ and the important contributions they make to Earth System functioning (Post 2020 CBD Targets 1 and 3). However, (semi-)natural habitats in human-modified lands and waters are often overlooked in conservation policies and global target setting despite the critical roles they play in supporting human well-being^3^ and conserving biodiversity^4^. Human-modified lands cover approximately 50% of the ice-free terrestrial land area ranging from urban areas to agriculture in mixed mosaic landscapes^5^. The loss of ecosystem function in such areas is incompatible with numerous sustainable development goals and targets discussed in the post 2020 Global Biodiversity Framework, notably target 10 on Sustainable Production^6^. Therefore, conservation efforts should address the functioning of human-modified ecosystems but lack specific metrics on the functional contributions that biodiversity embedded in human-modified lands make to support human well-being^7,8^.

Identifying such metrics is challenged by the highly context specific conditions under which biodiversity supports ecosystem processes in these human-modified ecosystems. Functional integrity has been proposed as a way to capture these diverse processes in a synthetic measure^9^ but clear evidence of the minimum level of functional integrity remains missing^10^. We define functional integrity as the capacity of the ecosystem to contribute to biosphere processes and to sustain multiple NCP through the presence of ecologically functional populations of species^9^. It addresses both Earth system-scale biosphere processes and provisioning of local NCP. Functional integrity differs from other biodiversity measures used in conservation biology as it recognizes that solutions can be provided by highly altered (non-native, or non-intact) functional ecological communities in agricultural, urban, and other human-modified areas.

The quantity, quality, and spatial configuration of (semi-)natural habitat (hereafter “habitat”) can be used as a proxy measure for functional integrity^11^. Habitat quantity refers to the proportion of (semi-)natural elements present in a landscape. Habitat quality is a measure of the ability of a habitat to host and maintain species required for specific ecological functions. The structure and composition of a habitat are strong determinants of quality^12^. The spatial configuration of habitat in the landscape influences landscape connectivity and the distribution of NCP providing organisms. The combination of quantity, quality and spatial configuration of habitat collectively underpin functional integrity.

The specific quantity, quality and spatial configuration requirements for NCP provision are strongly context dependent^13–15^. For NCP provided by non-mobile or low mobility functional groups (e.g. soil protection, capture of non-point source pollutants from surface and subsurface water, natural hazards mitigation), habitat location is extremely important. Sediment and nutrient capture are significantly improved through vegetation buffers along both sides of waterways, in particular stream headwaters^16^. Well-configured habitat significantly reduces the frequency and risk of natural hazards such as shallow landslides, floods, and soil erosion.

Numerous ecological studies have studied aspects of the relationship between habitat quantity, quality, and spatial configuration and the provisioning of NCP. Although these studies confirm the high context specificity and variability of such relationships, they consistently indicate that below thresholds of habitat quantity, quality, and spatial configuration, NCP provisioning is lost^17–21^. Pollination or pest control studies suggest thresholds of 10-20% habitat per km^2^ often based on expert judgement and valid for a specific management intensity level or landscape type^11,22^. However, a synthesis of minimum thresholds for functional integrity across several NCP and across a wider range of landscapes has not been conducted to date.

We conducted a systematic review of 153 peer-reviewed studies including 72 quantitative reviews and 81 narrative studies, comprising a total of 4458 original studies (see Supplemental Information) to assess the minimum level of functional integrity needed for NCP provision. We identify how much habitat is needed, of what quality, and in what spatial configuration, to ensure a minimum level of six critical NCP including (1) pollination, (2) pest and disease control, (3) water quality regulation, (4) soil protection, (5) natural hazards mitigation, and (6) physical and psychological experiences (hereafter “recreation”).

## Results

We estimated the minimum threshold of habitat quantity, quality and spatial configuration for the six aforementioned NCP. While local context is critical, we find a consistent pattern across studies showing that species, and the services they provide, are lost below a certain threshold of habitat quantity, and by the maximum linear distance that providers organisms can move.

### Habitat quantity

We find that at least 20% habitat is needed to support pollination and pest and disease control. Depending on context this minimum area may range from 10 to 50% for pollination, while it ranges from 10 to 38% for pest and disease control. For recreation provided by green spaces in and around urban areas at least 25% habitat (ranging between 19-30% depending on the context) is recommended (Figure 1, Table 1).

**Figure 1.**
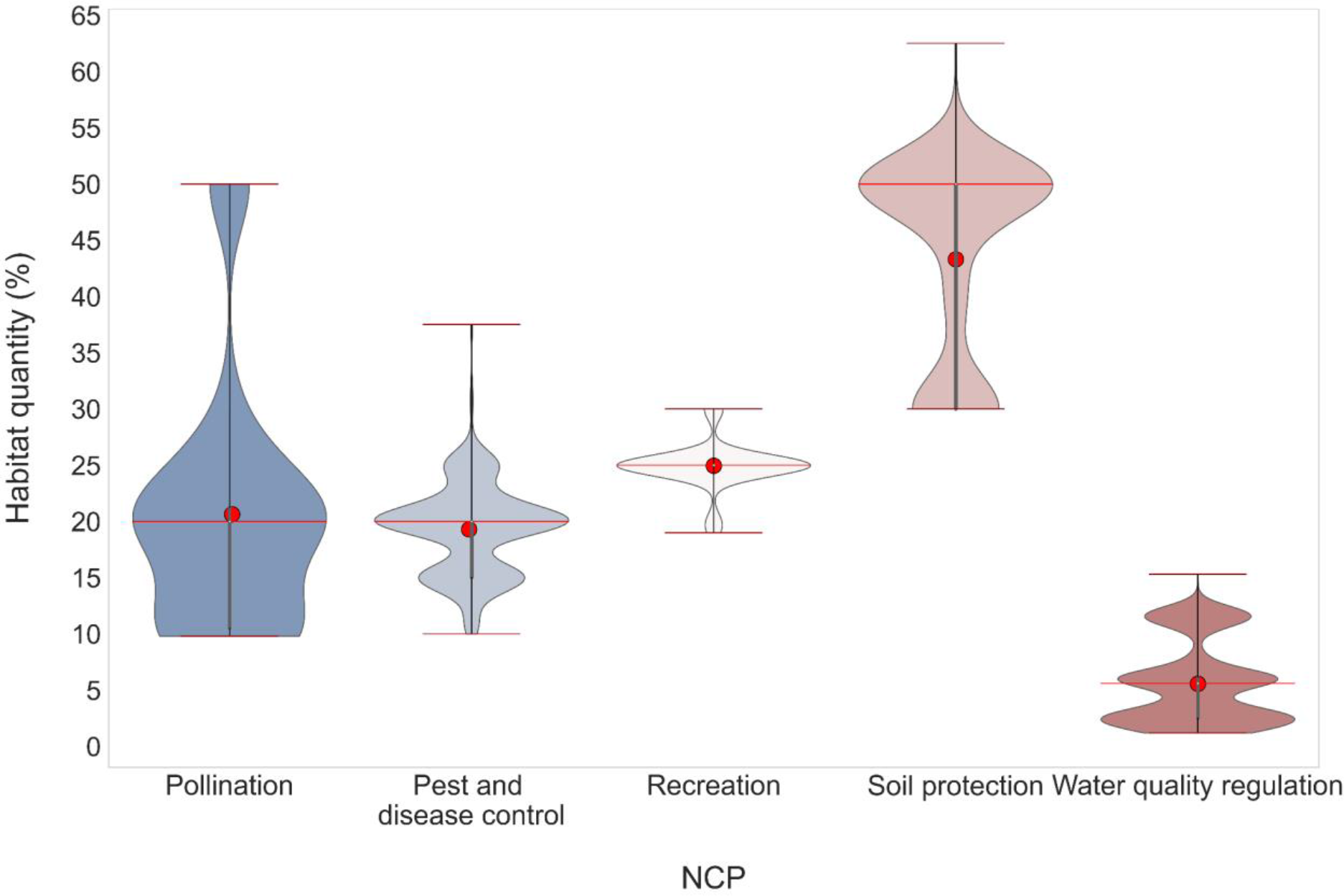
Threshold minimum quantity of habitat required for provisioning of nature’s contributions to people (NCP). The lower and upper red lines correspond to the whiskers (min, max, respectively) that indicate the range of the data.The middle red line represents the median, while the red dot represents the weighted mean value. The violin shape indicates kernel density estimation based on the number of original papers included in the (meta-)analysis reporting the given value.

**Table 1.** Threshold estimates for the major local ecosystem functions. The most constraining function (habitat quantity, quality or spatial configuration) per km^2^ is used to describe the threshold. All the values are weighted by the number of studies included. The total number of studies refers to the total number of studies considered in articles, reviews and meta-analysis.

| NCP | Taxonomic groups cited | Minimum median habitat quantity (%/km<sup>2</sup>) | Maximum median distance (m)/or position | Landscapes elements needed |
| --- | --- | --- | --- | --- |
| Pollination | insects | 20% (mean: 20.63±0.86 %) [10-50] (total 172 studies) | <500 m (mean: 843±32.3m) [15-2000m] (total 279 studies) | Rich diverse habitat with a diversity of native and non-native species (floral strips, floral field margins, floral under story cover; field grassy and woody margins, hedgerows, woody or silvo arable corridors between fields; forest edges and patches surrounding, grassland and shrublands patches). |
| Pest and disease control | insects, birds, arachnids | 20% (mean: 19.30±0.24%) [10-37.5] (total 260 studies) | <1000 m (mean: 760m±51.12 m) [10-2000m] (total 207 studies) | Complex habitat with diverse range of rich native species (forest edges and patches; floral strips, floral field margins, floral under story cover; grassland, pasture and shrubland patches surrounding; floral grassy and woody hedgerows and field margins; woody corridors between fields with floral understory) |
| Recreation | plants, birds | 25% (mean: 24.94±0.30%) [19-30] (total 50 studies) | <300 m (mean: 311m±6.7m) [300-500m] (total 44 studies) | Diverse rich semi-natural green spaces (streets trees canopy cover, public parks, zoos, gardens, woody and grassy parks, meadows) |
| Soil protection | plants | 50% (mean: $43.63 \pm 0.56\%$) [30-62.5] (total 251 studies) | Evenly distributed at the landscape scale | Diverse rich semi-natural vegetation cover (zoned grassy and woody buffers; trees canopy cover; ground cover with dense fibrous roots plants and cover crops such as grasses and legumes; agroforestry and woody and grassy hedgerows; mixed forest, shrublands and grasslands cover; extensive vegetation management with inter-row cover or crop cover, no-till farming, organic farms) |
| Water quality regulation | plants | 5% (mean: $5.6 \pm 0.09\%$) [1.2-15.3] (total 1480 studies) | both sides of streams | Native diverse semi-natural vegetative buffers or strips with diverse range of native species (three zoned buffers “native forest, shrubs and grasses”; forested or mixed forested and grassy buffers; grassy buffers or mixed buffers; wetland) |
| Natural hazards mitigation | plants | 50% (mean: 50.5%) (total 2 studies) | landslides: toe slope or slope bottoms | Semi-natural vegetation cover with diverse native species (native strong deep-rooted trees and shrubs with more reinforcing effect and low surcharge (low height and low diameter); spaced young exotic species (18-20 m) such as popular and willows; natural young trees; mixed plantation) |

The required quantity of habitat in the landscape needed to protect soil from water-based erosion is 50% (ranging from 30 to 63% for specific contexts, e.g. slope angle, precipitation intensity or landscape type) vegetation cover at the landscape scale, while for regulating streams water quality from non-point source pollutants requires about 6% habitat (a buffer of approximately 28 m width on both sides of streams). Total quantity of habitat needed ranges between 1.2-15% depending on the function in question, slope angle and stream density. Identifying the quantity of habitat for reducing landslide risk (natural hazards mitigation) is more challenging, with environmental variables (geology, slope geometry, soils, precipitation event frequency, intensity, and duration) often over-riding biological ones (vegetation quality) (Figure 1, Table 1). We found only two studies proposing a quantitative threshold limit for regulating landslide risk, advising a minimum of 50% and 60% vegetative cover on steeply sloped lands (>35°)^23,24^.

### Habitat quality

The capacity of a given area to generate multiple NCP is not only governed by habitat quantity but also by the composition and the type of habitats it contains. NCP are provided by communities of species and their traits. Vegetation characteristics define habitat quality and provide biophysical contributions such as sediment interception, which also serves as habitat for mobile species providing pollination and pest control amongst others. Our review identified six categories that we term ‘landscape elements’ for improving NCP provision: (i) complex diverse (semi)-natural habitat (SNH), (ii) complex diverse natural habitat (NH), (iii) diverse floral resources, (iv) forest, grassy elements and (vi) woody elements (Figure 2, SI Table 1). These landscape elements can be found in various forms within landscapes, including strips, patches, hedgerows, field margins, field borders, ground cover, canopy cover and buffers, and in urban areas, gardens, zoos and parks. Required habitat quality is NCP dependent with differences in specific needs (Figure 2). Nevertheless, 79% of studies reviewed found heterogeneous landscapes consisting of complex diverse (semi-)natural habitat is the most suitable for supporting multiple NCP provision (Figure 2). Increasing habitat floristic complexity and richness by including wild and native species and decreasing frequency or degree of disturbance (tilling, pruning, moving) strongly impact pollinating organisms diversity. Pest and disease control organisms are more diverse in complex diverse (semi-)natural habitats dominated by diverse woody or grassy elements rather than being determined by floral resources availability.

**Figure 2.**
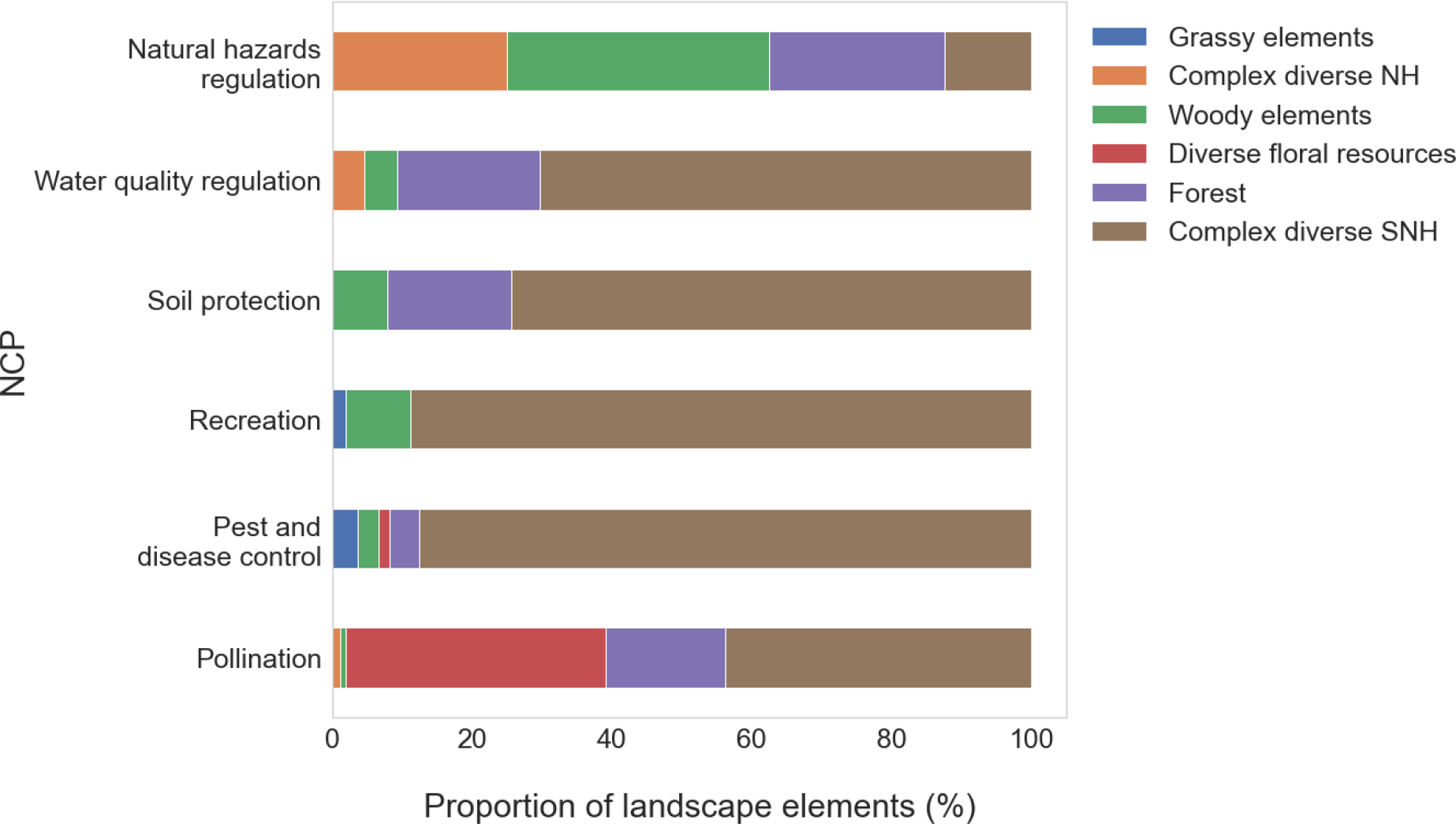
Landscape elements required for provisioning of each NCP. The stacked bar chart showing the proportion of landscape elements categories that improve habitat quality and support the provisioning of each NCP. Each bar in the chart represents a whole number of papers studied for each NCP, and segments in the bar with different colours represent different landscape elements categories. All the values are weighted by the number of papers.

The benefits from recreation spaces obtained in urban ecosystems arise mostly from structurally complex diverse (semi-)natural vegetation cover including street trees and public parks and green areas.

Preventing soil particle detachment from waterborne erosion depends on canopy and ground cover interception to protect soil from different types of water erosion (rill, gully, splash, or stream bank erosion). Preventing such erosion requires a structurally complex diverse (semi-)natural vegetation cover (including vegetated buffers, woody and grassy hedgerows or agroforests, ground cover or understory vegetation, inter-row vegetated strips, or crop cover with grasses or legumes) with a dominance of forest and woody elements.

Riparian buffers consisting of structurally complex, high species diversity plantings with a dominance of native species can be an important means of intercepting detached soil particles (sediment), pesticides and nutrients from adjacent fields. Important attributes include diversity of root structures, including both fibrous and tap rooted species to increase soil porosity, infiltration and excess nutrient capture. Slowing excess water flows through dense vegetation (e.g. zoned buffers consisting of grassy, shrub and woody elements) allows larger particles to fall out of solution and be retained in soils. On steep slopes, deep rooted perennial native plant cover from diversified fast growing plantings and understory vegetation are most effective to reduce landslides.

### Spatial configuration

Adjacency is an important element of NCP provisioning, notably in managed lands where distances between habitat (NCP source), and crops (NCP beneficiary) can vary.

Location of habitat within a landscape plays an important role in water quality regulation (riparian zones), whereas distance to source habitat determines access by NCP providing mobile organisms in the case of pollination, and pest and disease control where the foraging range from their home habitat determines maximum linear distance for NCP provision. This also applies to recreation where physical access or the reasonable distance that humans move to access green space determines potential use. For pollination and pest and disease control, notably insects, we identified a maximum linear distance of 500-1000 m (ranging between <0-2000 m for specific taxa) that insects can forage from their host habitat to the target crop field (Figure 3). For recreation obtained in urban ecosystems most studies indicate 300 m as a maximum reasonable distance for people to access green spaces based on the identified positive health impacts of experiencing at least >120 minutes of nature exposure per week^25,26^ (Figure 3). Beyond this distance the NCP provision declines significantly or is completely lost (Figure 3, Table 1).

**Figure 3.**
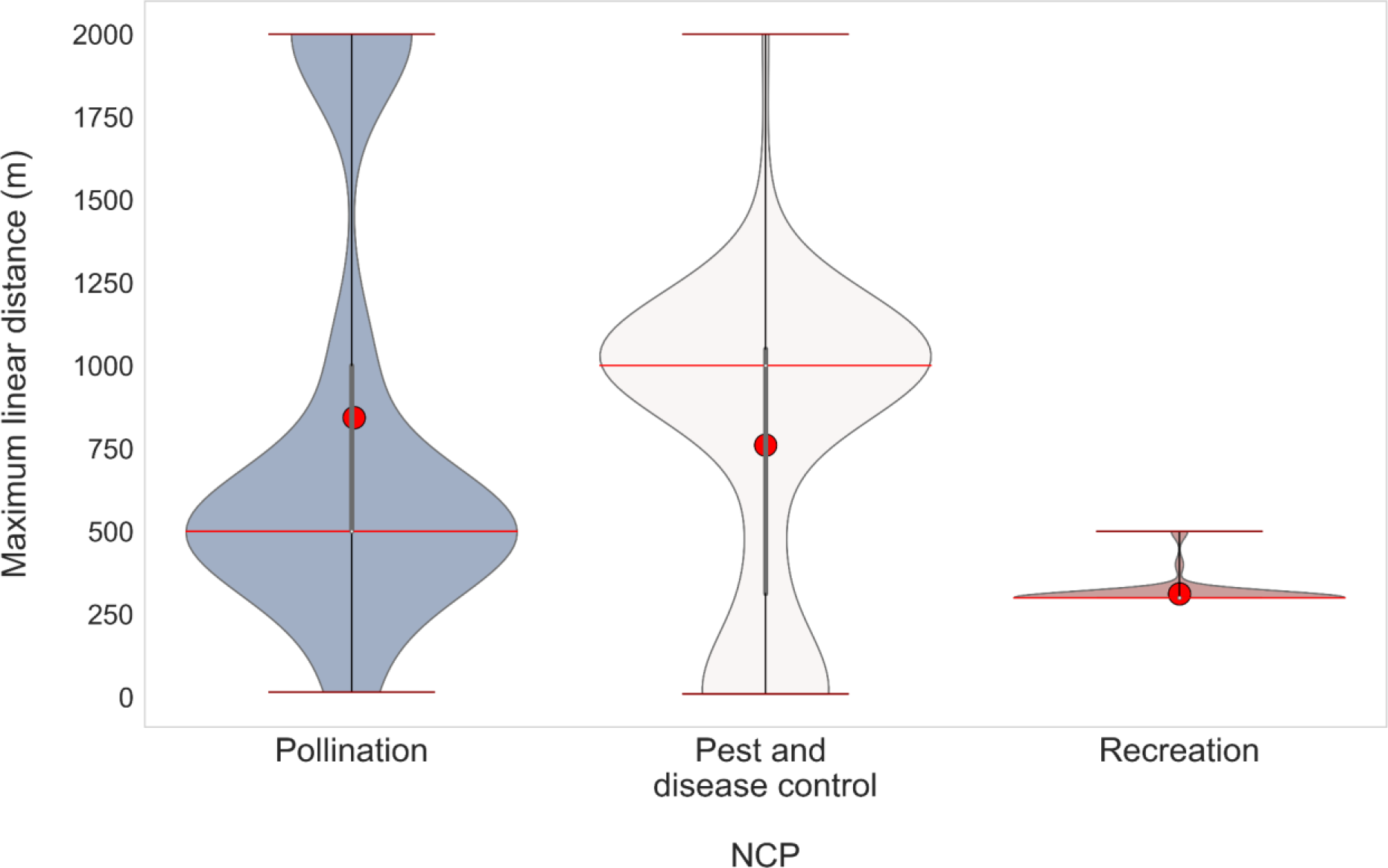
Threshold maximum linear distance values range (m) for each NCP: The lower redline and the top redline correspond to the whiskers (min, max, respectively) that indicate the range of the data, while the redline within the low redline and the top redline represents the median. The violin shape indicates kernel density estimation that shows the distribution of the values. Wider sections of the violin plot represent a higher probability that the number of the papers will take on the given value; the skinnier sections represent a lower probability. The red circles represent NCP’s mean maximum distances (m). All the values are weighted by the number of papers.

Reducing soil loss from waterborne erosion requires 50% of complex diverse vegetation cover evenly distributed evenly across the landscape on and around agricultural fields, and in uplands to reduce soil loss, on average by at least >71% (while ranging between 50 and 93% in specific contexts) (Table 1, Figure SI 1). The high minimum value for this contribution is driven by the mechanics of soil particle detachment, soil covering vegetation or litter.

For riparian buffers the spatial configuration needs are rather specific and concentrated around the streams. In spite of variation depending on the levels of pollutants and sediment, vegetative buffers of at least 28 m width on both sides of a stream headwater, close to the water body, notably on slopes <23° (on average) are generally able to capture >73% (ranging between 50-90% depending on the context) of nonpoint source pollutants(Table 1, Figure SI 1). This includes sediment, nutrients, pesticides, and salts from upstream agricultural lands. Considering global stream densities^27^, this buffer weight would on average, correspond to 6% habitat quantity per square kilometre.

As result of relatively sparse evidence, we did not identify a maximum distance measure for enhancing slope stability and reducing landslide occurrence on steep terrains (slopes >35°). Nevertheless, some studies indicate that on these slopes, retaining at least 50% complex diverse (semi-)natural vegetation cover distributed evenly with trees (the heaviest elements) placed mainly on the toe or the bottom of the slope is most effective^23,24,28^.

### Functional integrity thresholds

Based on our review of threshold levels for habitat quantity, quality and spatial configuration across the six NCP assessed, we propose a simplified, integrative, measure of functional integrity to secure minimum levels of NCP provision across global human-modified landscapes. We emphasize that our methodology focuses on minimum levels, and that increasing habitat quality, quantity, and proximity through targeted spatial planning can improve or increase NCP provisioning and efficiency. For habitat quantity, at least 20-25% habitat is needed to ensure that multiple NCP are provided (Figure 1). The quality of habitat retained is equally important, we find that landscapes consisting of complex diverse (semi-)natural elements are the most effective habitat across multiple NCP (Figure 2). In terms of spatial configuration, we identify 300 to 500 m as the maximum threshold distance between habitat and target beneficiaries (crops, citizens) for three of the six NCP evaluated: pollination, pest and disease control, and recreation (Figure 3). When these three are combined, we estimate that at least 20-25% complex diverse (semi-)natural habitat is required within each km^2^ of the landscape (Table 1). In landscapes with high erosion or landslide risk, a greater habitat fraction is needed. While some NCP may still be provided with habitat levels as low as 10-20%, in 90% of the studies based on habitat quantity we reviewed; the NCP provisioning is absent below 10% habitat.

### Current state and spatial distribution

Using the ESA World cover 10 m resolution land cover map of freely available satellite-based land cover data, we estimated the current state and spatial distribution of functional integrity by calculating the percent habitat in 1 km^2^ neighbourhoods, after distinguishing pasture land from (semi-)natural grasslands and testing for distinguishing forest plantations from (semi-)natural forests^29^. Our results indicate that 50% of human-modified lands, which account for 35% of all lands globally, are below 10% habitat per km^2^ and 64% to 69% of human-modified lands are below 20-25% habitat per km^2^, respectively (Figure 4, SI Table 2). This is significantly higher than previous estimates using lower resolution imagery^9^. While the limited thematic resolution of the land cover data may lead to an underestimation of habitat in the landscape (i.e. floral resources, grassy patches), it is, nevertheless, likely that about half of global human-modified landscapes are below the 20% per km^2^ habitat required to provide essential NCP, and are thus relying on substitutes for those NCP (domesticated honey bees, pesticides, technical means of water regulation and purification), or face absolute shortages in NCP. This shortage is especially found in the intensively farmed regions important to global food systems, threatening the long term resilience and adaptive capacity of food production systems.

**Figure 4.**
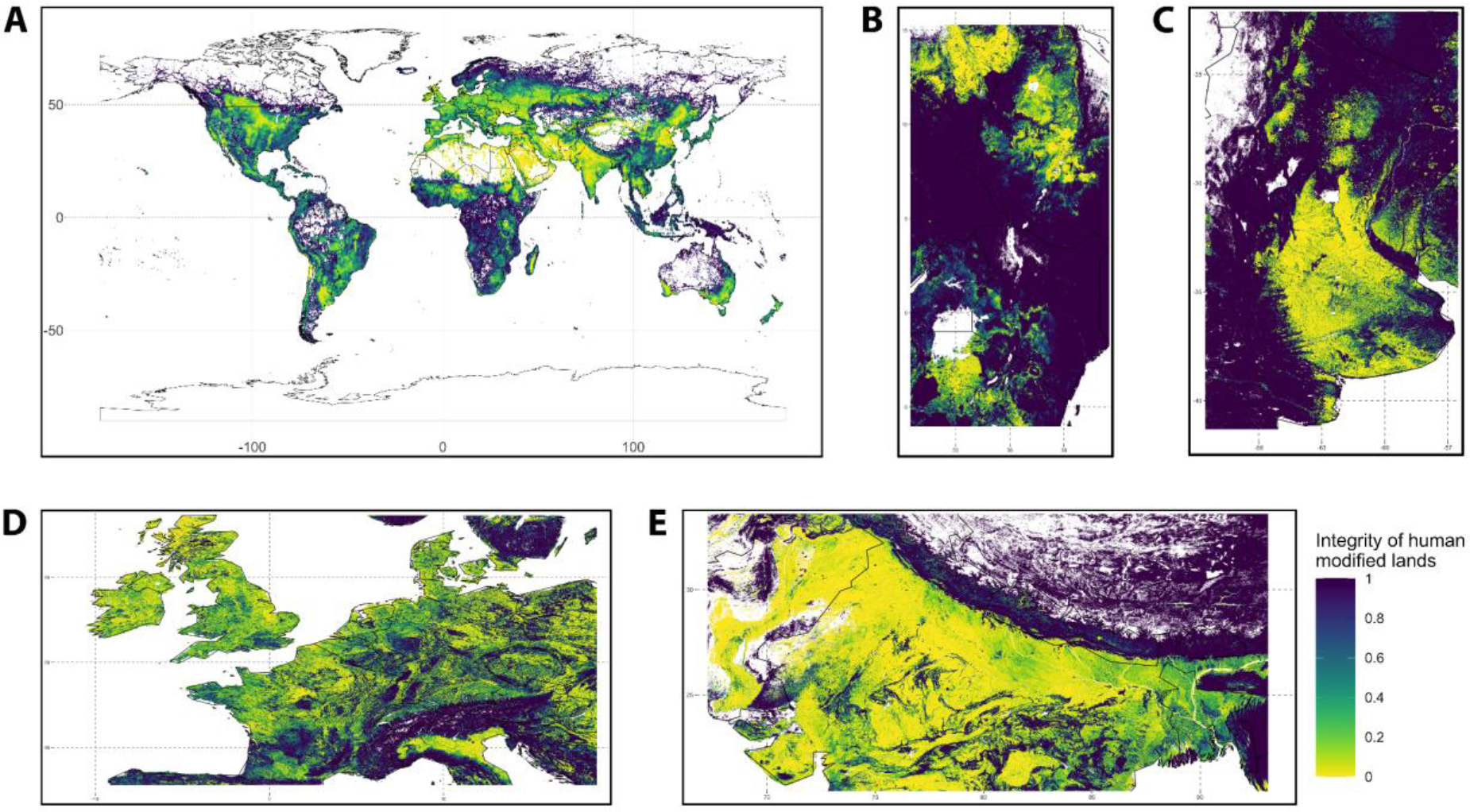
Functional integrity in human-modified lands. Per cell habitat integrity in human-modified lands (agricultural and urban landscapes) calculated as the percentage (%) of semi-natural habitat within 1 km^2^. Integrity is calculated at a 10m resolution and then aggregated for display purposes. (A) The global spatial distribution of biosphere functional integrity at a 500 metre scale. More detailed views are shown in the zoom-in panels at a 100 metre resolution for (B) East-African highlands and savannah, (C) Argentinian soybean region, (D) west-central Europe, and (E) Indian Gangetic plain. Areas coloured white indicate regions where there are no human-modified lands.

## Discussion

Our results, based on structured review of empirical evidence, propose that at least 20-25% complex diverse (semi-)natural habitat in each km^2^ of human-modified lands (Figure 1) is needed to maintain multiple NCP simultaneously. This requirement covers the quantity, quality and spatial configuration of habitat needed to minimally secure these NCP. We focused on six NCP and thus do not fully capture all facets of biodiversity and NCP that supports the needs of people. However, the selected NCP are representative of the fundamental principles by which biodiversity provides functions and contributions to human well-being. The functional integrity threshold we identified does not indicate the amount of NCP produced but rather indicates a threshold below which the provisioning of the NCP falls below a minimal level. When habitat quantity decreases further to a level below 10% complex diverse (semi-)natural habitat in each km^2^ of human-modified lands, five of the studied NCP are no longer provided. The minimum threshold is applicable to most human-modified landscapes^11,17,30^, though more habitat quantity may be needed in specific contexts, such as erosion-sensitive lands (see results section). Further, increasing landscape complexity through embedded habitat can further secure actual and future provision of NCP to humanity^31^.

Functional integrity as operationalized in this study is a useful measure as it can be captured with remote sensing, but it remains incomplete and biased to the role of above-ground habitats in securing NCP. Soil biodiversity contributions to soil quality, below-ground carbon sequestration, nutrient cycling and increasing water holding capacity in fields through no-till, or reduced tillage practices, cover crops or leguminous rotation are not captured by our measure despite their important role in improving soil health and function. Similarly, practices that reduce excess nutrient run-off are equally important and complement, but do not replace the role of habitat in buffering soil, nutrient and pollutants’ loss to aquatic ecosystems while supporting instream biodiversity^32^. However, excessive nutrient use can rapidly exceed the absorption capacity of riparian and other vegetated buffers, therefore reducing the pressures from human-modified lands increases the capacity of habitat to provide functional integrity.

Our analysis focuses on the value of increasing the diversity of habitat types in a landscape through the incorporation of complex diverse (semi-)natural elements (Table 1, Table SI 1). Complementary to this, increasing intra-field diversity of the agricultural/modified elements^21,33^, field edge density^17^ and decreasing field sizes^21^ may also increase landscape heterogeneity (but does not replace the positive effect of (semi-)natural elements on functional integrity). Which types of habitats are most appropriate to ensure functional integrity remains a highly local issue and should be driven by local knowledge.

Our findings show the importance of ensuring access to appropriate habitat (or landscape elements) at a sub-kilometre scale across human-modified landscapes. This is driven by the majority of species providing NCP having small home ranges, or being non-mobile. Numerous ecological studies show non-linear decreases in species diversity and abundance with increasing distance from habitat edges^34,35^. An additional benefit of embedding habitat within human-modified landscapes is to fragment agricultural lands. This reduces the dispersal of agricultural pests between fields^36^ while connecting habitat^37^. Securing riparian buffers is a good first step and would for example secure 6% of habitat per km^2^ on average globally while contributing to connectivity^38^.

The threshold of functional integrity identified from empirical data provides a useful measure for aligning global action. It emphasises contributions of biodiversity in supporting local NCP, for example those that either improve food production, or that reduce the negative impacts of food production. It provides a scaffolding of how much may be needed, which then requires local knowledge and locally adapted practices for implementation. Which practices are most suited to provide the six NCP analyzed here are best determined *in situ* and can span a broad range of practices cited in our systematic review (Table 1 SI, see Supplemental Information). Local studies and application, co-designed and conducted with local communities, will be critically important to identify the most appropriate interventions, and validate effectiveness^39^.

Historically, global monitoring of functional integrity of human-modified landscapes has been difficult as habitat mostly comes in small patches, often of linear format, that are not easily detectable in most coarse resolution global land cover maps. The recent high-resolution Sentinel images (10 meter resolution) used here are capable of capturing small patches of habitat as well as (most of) treelines and other landscape elements. However, these data might still underestimate habitat as they do not capture all hedgerows, field margins, floral strips and grass strips that are managed as (semi-)natural habitat. This is partially due to their limited spatial resolution, and also a result of the limited thematic resolution of this data-product. Unmanaged patches of grassland are not sufficiently distinguished by the data we used to distinguish grassland areas into pasture and throughout (semi-)natural grasslands. Similar concerns hold for forest land cover through remaining challenges of distinguishing natural forests from monocultures of short-rotation species. The sensitivity to distinguishing forest plantations from other forests was not large for the global results, but did show clear regional deviations (see SI, section 3.2). Given these limitations, our assessment of the current state of functional integrity should be interpreted with caution but remains useful to identify regions where functional integrity is likely to be below a safe threshold. We anticipate that with continued rapid evolution of remote sensing products and artificial intelligence these detention challenges are likely resolved in the near future. Early data have been published^40^, but are not yet openly available for inclusion in our assessment.

Despite these challenges, we estimate that at least half of the world’s human-modified lands fall below critical thresholds for functional integrity, severely compromising the capacity of human-modified lands to contribute to NCP provision. Restoring habitat in these places might compete with or complement increasing food production depending on the location and practices. For example, restoring habitat in agricultural lands does not necessarily result in reduced yields, with evidence that a diversity of practices both improve yields and environmental outcomes, and that embedded biodiversity on field perimeters and riparian buffers leaves scope for sustainable intensification within fields^41,42^. Other benefits of habitat in human-modified landscapes may include: a 10% increase in tree cover in agricultural landscapes may make a significant contribution to global carbon sequestration^43^, and small patches of habitat in human-modified landscapes may have disproportionate value in preserving species diversity^44^. The generalized trade-off between the area of natural habitat and food production is thus a false one resolved with locally appropriate conservation options. Innovation, notably in agroecological practices, can help to integrate new habitats in these landscapes. Restoring habitat and the ecosystem functions in human-modified landscapes can strengthen the resilience of ecosystems and contribute to halting the decline of biodiversity, as well as supporting the wellbeing of people by facilitating their access to nature’s benefits.

## Methods

### NCP selection

We selected NCP which are defined by clear ecological processes, notably regulating and supporting NCP. These include (1) pollination, (2) pest and disease control, (3) recreation, (4) soil protection, (5) water quality regulation, and (6)natural hazards regulation. Using a systematic approach in accordance with the Preferred Reporting Items for Systematic Review and Meta-Analyses guidelines [PRISMA] (Figure SI 2, see Supplemental Information for additional detail and references included), we searched for literature that addressed three key variables for each NCP to describe the minimum level of functional integrity that secures the ecosystem function underlying the provision. First, a quantitative measure of the minimal area of habitat needed to provide the NCP. Second, a qualitative evaluation of the type and quality of the landscape elements required, and third, the maximum distance between providers and beneficiaries (m) or the spatial configuration of landscape elements required for the NCP to be provided. The search yielded a total of 153 articles, comprising 72 meta-analysis and review papers and 81 primary research articles (Figure SI 2). We performed exploratory analyses to identify generalizable patterns in the studies regarding each NCP in question.

### Minimum values range calculation

The minimum threshold under which NCP are no longer delivered has been estimated and extracted either directly from papers’ text, tables or supplementary information or from the figures. In the figures’ case, we estimated the minimum threshold of habitat quantity for pollination and pest and disease control when either the abundance or diversity of NCP providers dropped significantly before crossing the zero or curve’s starting point value (Figure SI 3). For recreation in urban ecosystems, the minimum amount of green space under different forms and quality, as well as its spatial configuration or linear distance (see Table 1) from each neighbourhood has been assessed from several studies. For example, these studies analysed the relationship between the amount of green space in each neighbourhood in cities and peoples’ mental and physical well-being, measured by psychological distress level, number of natural-cause mortality, cortisol levels, prescriptions for antidepressants, presence of anxiety, COVID-19 incidence rate and heat stress level^45–47^.

For soil protection and water quality, the soil loss reduction efficiency of vegetative and/or buffer effectiveness or pollutants reduction capacity, respectively, have been considered as a baseline to estimate the minimum vegetation cover and buffer width needed to maintain the provisioning of the NCP. However, the reduction efficiency of vegetation buffers or cover is highly variable across studies and dependent on the NCP and landscape type, with no suitable reduction efficiency proposed across the studies. In 90% of the studies reviewed, the reduction efficiency of different amounts of vegetated buffers exceeds >50%. In our analysis we used >50% buffer or cover effectiveness or reduction rate as a baseline to determine the minimum value required. The buffer width is represented in meters; we then transformed the buffer width into an approximate amount of (semi-)natural vegetation using the average density of streams globally^27^.

For landslide mitigation, the minimum value has been determined from several experimental and modelling studies that calculate the factor of safety (FoS) with presence and absence of plant roots in the soil^48–51^. The factor of safety (FoS) is a crucial indicator of slope stability and is defined as the ratio of the resisting force to the driving force along a failure surface^51^. To have a stable slope, the FoS threshold of 1.3 is often specified for temporary or low risk slopes and 1.5 for permanent slopes^52^. Thus, we use the 1.3 FoS as a baseline and proxy to determine the minimum vegetation cover needed for maintaining slope stability.

We described habitat quality through the landscape elements recommended as suitable by the studies we reviewed for the survival of individuals and persistence of populations that provide these NCP. We identified six categories of landscape elements encompassing all of the studies reviewed: complex diverse (semi-)natural habitat (SNH), complex diverse natural habitat (NH), diverse floral resources, forest, grassy elements and woody elements (SI Table 1, see SI methods for more details).

For spatial configuration estimation, for three of the six NCP that are provided by mobile organisms (pollination, pest and disease control, and recreation), we extracted and estimated the maximum linear distance that mobile organisms forage or move to get access from their home habitat using the same approach as habitat quantity estimation. For the other three NCP that are provided by non mobile organisms (water quality regulation, soil protection, and natural hazards regulation), recommended emplacement and location of habitat providing the NCP has been extracted from the text of the relevant studies.

Once the thresholds of each NCP were determined at landscape scale, we analyzed what characteristics of functional integrity (habitat quantity, habitat elements and spatial configuration that are essential for functioning) are important for decision-makers and management.

### Functional integrity – current state and spatial distribution

We calculated the current state of the functional integrity boundary based on the ESA Worldcover 10 meter resolution land cover map (https://esa-worldcover.org/en), refining the grassland category by distinguishing pasture lands and (semi-)natural grasslands using the habitat map of Jung et al.^29^. We reclassified this to create a binary classification of “natural lands” and “human-modified lands”. We then calculate an integrity value for each pixel using a focal function where we calculate the mean of the binary for the 500-metre radius around each pixel and calculate the percentage of pixels that meet different thresholds (10%, 20%, 30%, etc.). We performed an additional sensitivity analysis using the Jung et al.^29^ classification to refine the ESA Worldcover ‘tree cover’ category by distinguishing forest (seen as natural) from plantations (seen as human-modified lands). For full details see the Supplementary Information.

## Supporting information

Supplementary NCP specific supporting information, supplementary methods, supplementary figures, supplementary tables and references

## Acknowledgements

This work is part of the Earth Commission which is hosted by Future Earth and is the science component of the Global Commons Alliance. The Global Commons Alliance is a sponsored project of Rockefeller Philanthropy Advisors, with support from Oak Foundation, MAVA, Porticus, Gordon and Betty Moore Foundation, Herlin Foundation and the Global Environment Facility. The Earth Commission is also supported by the Global Challenges Foundation. DeClerck received additional support from the Food System Economics Commission and Verburg from the NatureConnect project funded by the European Commission.

## Data Availability Statement

All data analysed in this study are available from the corresponding author on request. Data that support the findings of this study are available within the paper and its references and Supplementary Information.

## Code Availability Statement

The code used to analyse the data and produce the figures and maps is available from the corresponding author upon request.

## Author contributions

AM developed the methodology for assessing and analysing functional integrity, participated in the conceptual design, conducted the systematic review, gathered and analysed data, led the write-up of the paper, and served as a research scientist on the Earth Commission’s Biosphere working group.

FD, PHV, DO originated the idea, developed the concept and methodology for assessing functional integrity, contributed to the analysis and write-up and co-led the Earth Commissions Biosphere Working Group.

JFA participated in the conceptual design and writing of the paper, performed the spatial integrity analysis, created the spatial maps, and served on the Earth Commissions Biosphere Working Group

NZC contributed to the analysis and the writing of physically and psychologically beneficial experiences in nature and served as a member of the Earth Commission’s Biosphere working group.

NE-C, AF and SJ contributed to the conceptualisation and methodology for assessing functional integrity and to reviewing of the final manuscript

JR participated in the conceptual design and writing of the paper and served on the Earth Commissions Biosphere Working Group.

ICM contributed to the analysis of the soil protection NCP and to reviewing the final manuscript.

BSK contributed to the riparian analysis and reviewing the final manuscript.

