## Supplementary NCP specific supporting information, supplementary methods, supplementary figures, supplementary tables and references for "Biosphere functional integrity for people and Planet"

**Including *Supplementary:***

1. ***Context dependent discussion supporting information: including mechanics of NCP provisioning***
2. ***Methods***
3. ***Figures***
4. ***Tables***
5. ***References***
6. ***List of references used in the NCP analysis.***

#### *1.1. Mechanics of NCP provisioning*

Substantial local variations are the norm in NCP research, notably in relation to the study of single species contribution to a single service. The exact quantity can vary strongly with landscape type[1], management regime intensity, agricultural practices and crop diversity[2–4], crop type[5], field size[3], field edge density and NCP providers or taxa[6]. For example, the quantity of habitat needed increases with increasing management intensity, however increasing the quantity of habitat (>20%) in such intensified landscapes may be difficult to achieve and has a negative impact on crop yield. The alternative solution can be to increase crop diversity (doubling at least) which has a positive effect on multi-trophic diversity (e.g. 4 times more pollinators) if the habitat quantity is at least maintained at >10-11%, supporting the "landscape complementation" hypothesis[3,7]. Moreover, reducing mean field size and crop-crop borders has been shown to increase multi-trophic diversity and enhance landscape connectivity even in the absence of habitat between fields[2,8–11]. The effect of habitat quantity is primarily impactful on arthropod diversity[7,12,13]compared to arthropod abundance which shows inconsistent responses, notably across specialist and generalist arthopods[14,15].

Our survey of the literature also found that maintaining at least 25% of complex diverse (semi)-natural vegetation cover in every neighbourhood in cities is needed to support people's mental and physical health. Many studies show the importance of increasing green spaces in cities to reduce psychological stress levels, cortisol levels, prescriptions for antidepressants, presence of anxiety and premature mortality, while contributing to the developing healthy, liveable and sustainable cities e.g. *Kondo et a*l.[16] have demonstrated that 400 deaths annually could be prevented when increasing the canopy cover from 20 to 30% of the city land surface, *Mueller et al*.[17] have shown that 60 deaths annually could be prevented in one neighbourhood when increasing the proportion of the green spaces from 6.5%-19.6%.

Fifty percent of complex diverse (semi)-natural or natural vegetation cover emerges as a minimum requisite to protect soil from water erosion and can reduce soil loss by more than 71%. However, countless studies on which this minimum quantity of habitat was based are conducted on erosion sensitive terrains, and most likely the quantity of vegetation required to prevent soil loss is much lower in less sensitive areas. Vegetation cover has an important impact on reducing soil loss and runoff via the redistribution of rainfall into three components – canopy interception, stemflow and throughfall – thus limiting runoff and controlling different mechanisms of soil erosion including rill and inter-rill, splash and gully erosion. Previous studies have proposed >40-60% vegetation cover as a critical threshold below which accelerated erosion dominates, notably on sloped terrains. Below the proposed threshold, the risk of soil erosion is extremely high (e.g. 100-1000% increase in erosion rates)[18–20].

Preventing surface and subsurface pollution from entering streams, lakes or wetland ecosystems from agricultural upland areas requires a minimum complex diverse riparian buffer threshold of 28 m protection on each side of the stream in question. Scaled globally, this equates to approximately 6% (1.2-15%) per km$^2$. As with other NCP this threshold varies depending on local context conditions including slope, topography and ecoregion climate regime, upland land-use management intensity, rainfall intensity, target water (surface or subsurface) and pollutant in question (e.g. nutrients, sediments, pesticides). For example, buffer width increases when increasing slope with the need of a mean addition of 0.8-1 m to the baseline buffer width for each 1% increase of slope[21]. Wider buffers are needed in tropics than in temperate ecoregions[22]. Greater buffer width is also needed when adjacent land use intensity is high or if the target objective of the management is to maintain biodiversity[23,24].

For landslides in hilly landscapes (>30°), although vegetation has extensively been used as natural protection against factors that trigger landslides and debris flow, only two studies (from China and New Zealand) proposed quantitative boundary limits of at least 50% of (semi)-natural vegetative cover)[25,26].

Although 20-25% of habitat secures the provisioning of multiple NCP, the spatial configuration, linear distance or location of habitat, as well as its quality all together must be considered in landscape management to enhance biodiversity, ensure ecosystem functional integrity to maintain desired levels of functions and services.

The implementation of our 20-25% threshold is applied to 1 km$^2$ resolution. The exact configuration and quality of habitat needed depends on the landscape type, management regime, topography, function in question and the functional groups that provide the NCP leaving important flexibility and choice options to local communities in identifying and applying the most appropriate practices backed up by available local ecological evidence.

We found that for mobile functional groups, notably insects that support food production services, complex diverse habitat including sufficient floral resources configured into different forms (e.g. floral strips within fields, floral, grassy and woody field margins, hedgerows, woody corridors, forest edges and forest, grassland and shrublands cover) is needed to support mobile arthropod diversity and maintain services provisioning. Habitat patches surroundings should be evenly configured into many evenly dispersed small patches, within a maximum linear distance of 500 m and 1000 m from the target crop field to support both pollination and pest and disease control services respectively. Beyond this distance the NCP in question is lost or no longer accessible. This is in agreement with many previous studies demonstrating the important positive impact of habitat proximity to the target crop notably for mobile biological functional groups[27,28]. The effect of the isolation varies highly across taxa and depends on the availability of complex diverse resources for nesting or matting sites, quality of habitat as well as the foraging ranges of each taxa[28–31]. For example, the isolation of habitat from the target crops (>1-2km) significantly reduces the visitation rate of bumblebees, hoverflies and solitary bees rather than impacting honey bees or butterflies who have longer distance foraging ranges and thus are less affected by isolation[31–33]. Both pollinators and natural enemies were also more diverse and abundant in complex diverse and species rich habitats than less diverse regardless of the form of habitat[31,34,35]. Moreover, pollinators and natural enemies respond distinctively to major habitat types with more diverse and abundant natural enemies along perennial habitat (e.g. hedgerows)[36] than herbaceous annual. Pollinators, in contrast, especially bees were in general higher in herbaceous or grasslands habitat with sufficient floral resources compared to woody habitat[37,38]. Sufficient continuous, unbroken hedgerows, field margins, between fields corridors, small, dispersed habitat and proximate forest edges surroundings, floral strips within farms, with a diverse range of native and rich species are optimistic strategies to maintain biodiversity and produce food in a sustainable way.

We also found that 25% of complex diverse green space within a linear distance of 300 m from every home in cities (equal to 5 min of walk from each home[39]) is needed to support mental and physical human health. Exercising or walking in nature results in lower cortisol level, lower prevalence of depression, anxiety and stress even under high periodic life stress compared to physical exercise alone or by watching nature scenes[40]. Green space accessibility and quality are among the strongest indicators driving green space's contributions to physical and community wellbeing[41]. Green space quality including plant, butterfly and bird richness has a positive effect on well-being[42,43], although in some studies the impact stays neutral. The spatial configuration of green space also has important feedback on human mental health with less psychological distress reported by residents within urban landscapes that have a disaggregated distribution of urban green spaces (i.e., many small green spaces, small-sized water bodies) compared to a single or large green space. The implementation of 25% complex diverse (semi-)natural vegetation cover dispersed into small green spaces within 300 m from each home in cities can contribute to wellbeing and the development of sustainable, liveable, and healthy cities.

For protecting soil from different types of water erosion and maintaining soil productivity, we found that a target of 50% of complex diverse mixed (semi-)natural vegetation cover under different forms (see results section) to be established and dispersed evenly across the landscape is needed. Soil erosion process results in the loss of the most fertile topsoil and accelerates because of human activity including land use and farming practices, as well as climate change. The efficiency of perennial vegetative cover in reducing soil loss depends on the vegetation type, slope, target erosion type, soil type and ecoregion[44]. Forests are more efficient in reducing soil loss in sloping farmlands (slope >25°) compared to grasslands or shrublands that are more effective on >0-25° slope, while croplands or orchards demonstrated the highest soil loss value among land-use types[44]. For reducing a splash erosion, establishing a community of shrubs with sufficient canopy cover is requested[45,46], whereas for rill and ephemeral gully erosion plant roots are as important as vegetation cover[47]. High vegetation density agroforestry systems can reduce soil erosion rates by 50% compared to crop monocultures[48]. Soil erosion rates can be significantly reduced if the vegetation cover type is carefully chosen depending on landscape type, management objective and slope.

For riparian buffers, we found that maintaining 28 m ( 6%) complex diverse vegetated buffers including native species on both sides of the streams reduce about >70% of non-point pollutants entering from upland agricultural fields. The buffers removal or reduction efficiency is largely controlled by buffers emplacement, vegetation type, plant density and buffer zone width[49,50]. Implementation of vegetated buffers on the headwater streams is vital to maintain water quality in streams. Grassy buffers or mixed grass-woody vegetation are more effective at trapping sediment than at removing total nitrogen[50], whereas forested buffers are more effective at removing excess nitrogen[51,52] or phosphorus from the surface runoff[49]. The quality of the vegetated buffer should be designed based on the management objective and local condition. Overall, consideration of the inclusion of a diversity of species, including but not restricted to native species, distributed in either two or three tiered (forested, grassed) or mixed buffers combined woody and herbaceous species or wetlands can be used to trap non-point pollutants and maintain physical, chemical integrity of the streams, lakes and wetlands. The first buffer zone is recommended to be placed along the stream bank and to be likely planted with a narrow native forested buffer and a second wider zone of native woody vegetation extended from the edge of the first zone through land and the third buffer zone should be narrow and extend upslope from the edge of the second zone. The 28 m width of a complex diverse (semi-)natural vegetated buffer should be applied on both sides of the headwater streams where the removal efficiency and processing is mostly achieved. Agricultural practices and urbanisation (e.g.roads) should be also carefully planned across whole catchments in order to maximise benefits[22].

Vegetation contributes to slope stability via two mechanisms including changes in soil moisture regime and contributing to soil retention by plant roots. Factor of safety (FoS) has been used in vast modelling studies as a proxy of vegetation cover effectiveness in reducing landslides and stabilizing slopes[53,54]. FoS is estimated based on vegetation transmissivity, soil physical properties, soil cohesion and additional cohesion from plant roots[55,56], Slope geometry, rainfall intensity, vegetation type, plants roots architecture and depth, the locationof trees along a slope, as well as their size, age and density, also impact the FoS[57–59]. For example, higher slope stability has been found in natural hillslope where native mixed forest species are introduced (FoS of 1.3) compared to modified hillslopes (e.g. hillslope with road cuts FoS of 0.95) despite a higher pore pressure in natural hillslopes[58]. FoS decreases with increasing rainfall intensity notably in modified hillslopes. Trees are more effective in reducing landslide occurrence by 95% than pasture or other vegetation types[59] when they are placed on the toe of slopes (e.g. increase the FoS by 7%), while positioning trees mid-slope or on ridge tops decreases FoS by 43% notably in hilly terrains (>35°)[46,60]. However, slopes >60° have a low probability of sliding if they have upwards of 80% of their entire slope length in perennial vegetative cover composed of mixed tree species dominated by natives[61]. Overall, to reduce landslides occurrence and increase the stability of hilly slopes and thus avoid harm to people, a suitable vegetation quality composed of mixed tree species spaced a maximum between 18-20 m along the toe of slopes should be considered to reduce risk of slides.

### 2. *Supplementary methods*

#### *2.1. Literature survey*

##### *2.1.1. Search strategy*

We performed a literature survey for reviews in accordance with the Preferred Reporting Items for Systematic Review and Meta-Analyses guidelines [PRISMA[62]] using main searches based on standardised two to three key-words and terms related to each NCP we analyzed:1) pollination ("pollinat*" AND "habitat" AND "landscape, 2) Pest and disease control ("Biological control*" AND "habitat" AND "landscape"), 3) physical and psychological experiences ("physical AND psychological*" AND "wellbeing*" AND "nature*"), 4) Water quality regulation ("riparian buffer*" AND "width*"), 5) Soil protection ("soil erosion*" AND "vegetation*" AND "landscape*") and 6) Natural hazards ("landslide*" AND "vegetation cover*"), in the 'Web of Science' (Clarivate Analytics, Philadelphia, USA). Additional papers including both articles and reviews papers were identified from other sources whether proposed by experts, manual search of the reference lists of all included reviews for additional reviews or by a similar search in google scholar engine using an additional search string for each NCP (e.g. 'pollination OR pollinators*'; habitat*; landscape configuration*; landscape complexity*; landscape heterogeneity*) between 2010 and 2021 and screened the titles and abstracts of the first two pages.

##### *2.1.2. Eligibility criteria*

Criteria for inclusion were defined in advance. We used the following guidelines for inclusion of potentially relevant references for the next steps of evaluation: Reviews had to be published in peer-reviewed journals between January 2010 and December 2021 in English. The source should focus on which taxonomic groups provide that service, or what food web connections are essential, quality of (semi-)natural habitat including landscape elements underlying the quality, distance or location or emplacement of landscape elements and some description of the spatial relationship between biodiversity and the NCP in question. There were no restrictions with respect to methods or landscape type or location of the study. All references not fulfilling any of the above conditions or being clearly out of scope (did not report outcomes on quantity, quality or spatial configuration) were excluded from the analysis.

##### *2.1.3. Study selection*

We first screened all papers (n= 411), initially based on titles and abstracts to identify potentially eligible reviews and relevant papers. We skimmed through the full-text articles to further evaluate the quality and eligibility of the studies. We assessed full papers with respect to the inclusion criterion. Papers not meeting the inclusion criteria were excluded. The flow of studies is presented in respect to the PRISMA diagram shown in Figure SI 2.

##### *2.1.4. Data extraction and management*

We extracted data from all eligible reviews and articles and tabulated them using a set of data extraction forms, which were developed for the present study. We collected the following information: name of the first author, year of publication, name of the journal, location of the

study, methods, nature of the paper, number of papers included, minimum habitat quantity, description of habitat elements, landscape elements recommended, maximum linear distance from source habitat or the location and emplacement of habitat, functional group providing the NCP, NCP providers responses to quantity, quality and spatial configuration (+, -, 0 or inconsistent), evidence and key results for each NCP according to the authors of the paper.
Habitat quantity, maximum distance and a description of landscape elements were searched and extracted either directly from papers' text, tables or supplementary information or from the figures. In the figures' case, we estimated the minimum threshold of quantity of habitat or maximum distance beyond which the NCP in question is lost when the value dropped significantly before crossing the zero or curve's starting point value (Figure SI 3 ). This value refers to different variables in interaction with habitat or quantity of nature needed depending on the NCP in question (see method section for more details).
For habitat quality, we grouped all potential landscape elements that improve habitat quality and are recommended in each paper into six groups that are: Diverse floral resources, complex diverse SNH, complex diverse NH, woody element, grassy elements and forest distributed or dispersed in different forms (see table SI 1 for more details).
We then performed exploratory analyses to identify generalizable patterns in this literature regarding the three key variables for each NCP in question using Python language (Python 3.6) in Anaconda (Seaborn and matplotlib packages).

#### *2.2. Functional integrity current state and spatial distribution calculation*

We calculated the current state of the functional integrity boundary using freely available satellite data, building on methods developed in DeClerck et al.[63] We used the ESA Worldcover 10 m resolution land cover map (https://esa-worldcover.org/en) to create a binary classification. We first reclassified the 'grassland' category to 'natural grassland' and 'pasture-land' by overlaying the habitat map from Jung et al.[64] Where the ESA Worldcover area that was classified as 'grassland' overlapped with the area classified as 'artificial – terrestrial' by Jung et al.[64] We reclassified this to 'pasture-land'. All other 'grassland' areas were reclassified as 'natural grassland'. We then converted this classified land use map into a binary classification where:

We calculated an integrity value for each pixel using a focal function where we took the mean of the binary for the 500-metre radius around each pixel. We then aggregated the percentage of pixels that meet or exceed different 'integrity thresholds' (10%, 20%, 30%, etc.) in both human-modified lands and all lands on a global scale and ecoregion scale.

We performed an additional sensitivity analysis using the Jung *et al.* classification to refine the ESA Worldcover 'tree cover' category. We reclassified pixels where the ESA Worldcover area classified as 'tree cover' overlapped with areas classified as 'plantations' by Jung et al.[72] as 'plantations'. All other ESA Worldcover 'tree cover' pixels were reclassified as 'natural tree cover'. 'Natural tree cover' was assigned a 1 and 'plantations' was assigned a 0 in the binary classification. We then followed the same procedure as above to calculate integrity. All analyses were done using the native Google Earth Engine interface.

### 3. *Supplementary Figures*

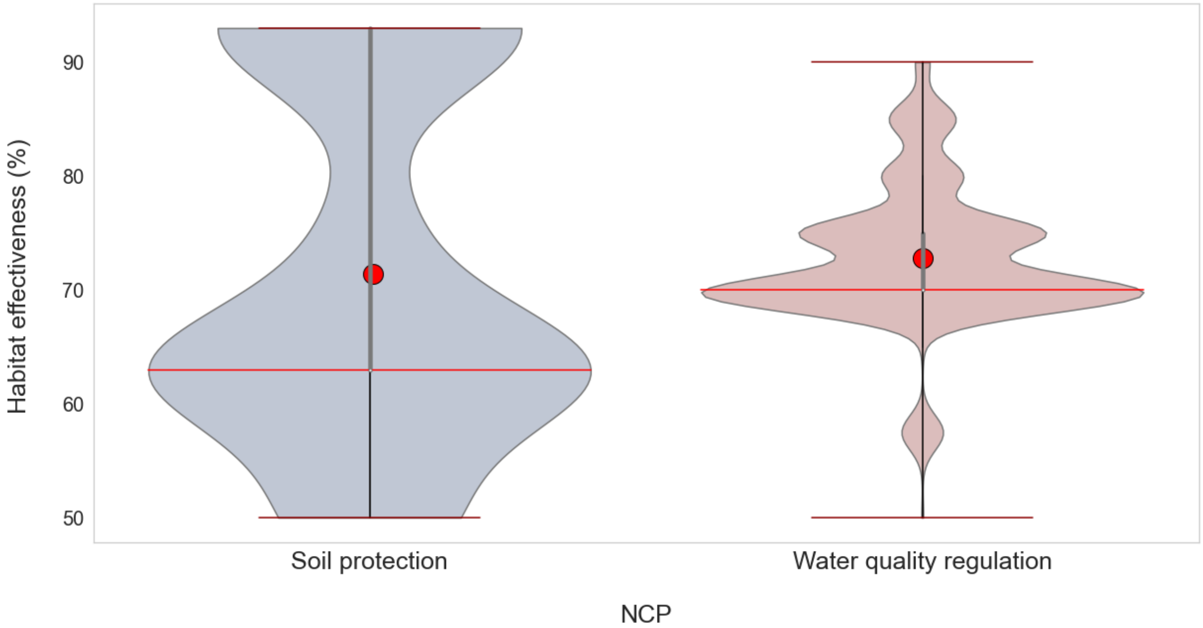


***Figure. SI 1. Habitat effectiveness threshold*** *(%) for soil protection from water erosion (gray) and water quality regulation from non-point pollution (light red). The lower redline and the top redline correspond to the whiskers (min, max, respectively) that indicate the range of the data, while the mid-figure redline represents the median. The violin shape indicates kernel density estimation that shows the distribution of the values. Wider sections of the violin plot represent a higher probability that the number of the papers will take on the given value; the skinnier sections represent a lower probability. The red circles represent NCP's mean habitat effectiveness (%). All the values are weighted by the number of papers.*

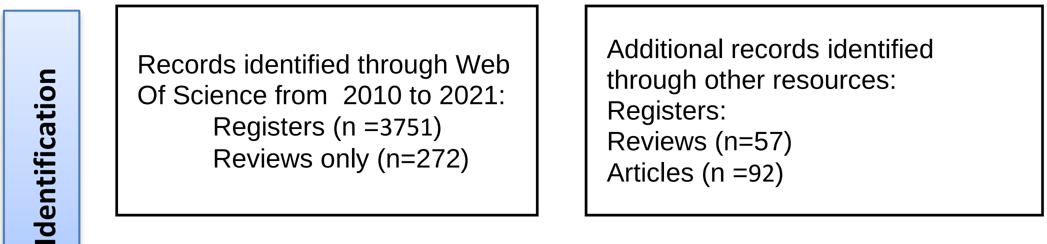

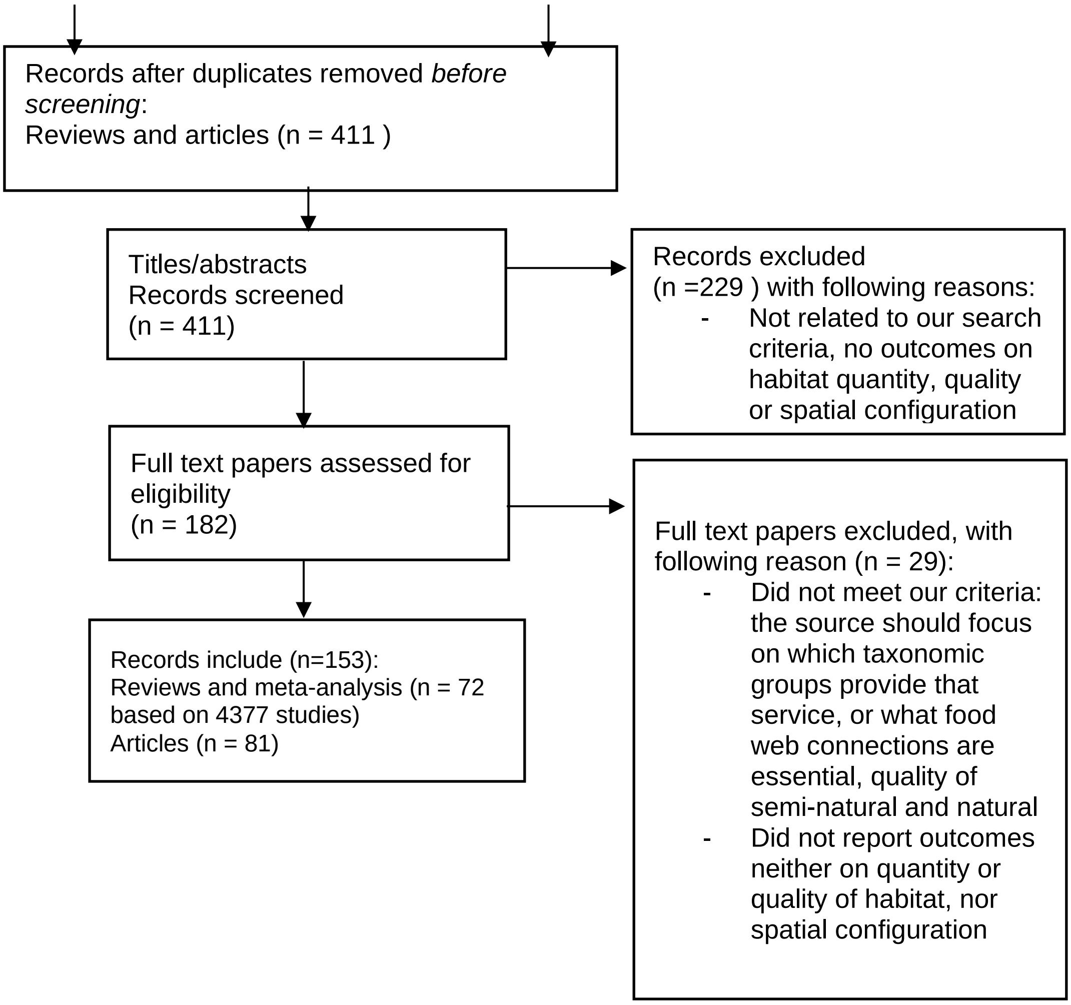


***Figure SI 2.*** ***PRISMA flow diagram for systematic reviews which included searches of databases and registers***

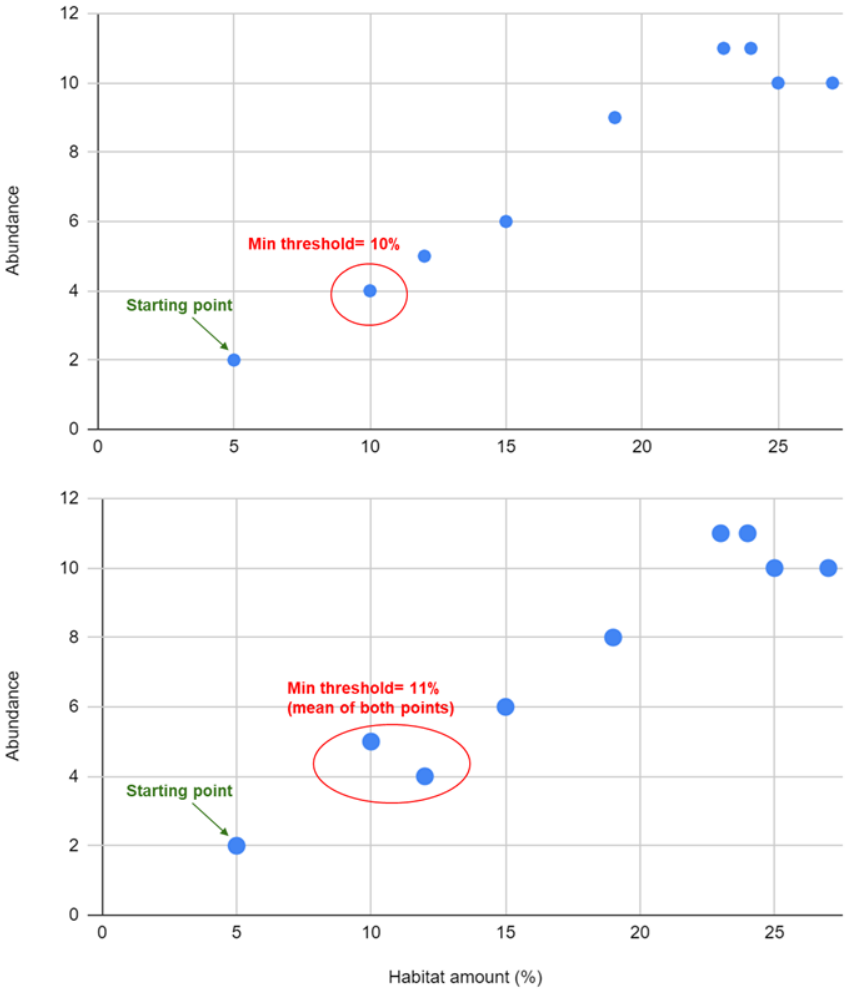


***Figure SI 3. Illustration figure*** *showing two examples of the method of extracting the minimum values from the reviewed studies figures when the starting point is not zero.*

### *4. . Supplementary Tables*

***Table 1. Landscapes elements categories and included elements underlying habitat quality for each category across 150 studies reviewed including 75 articles and 75 review and meta-analysis papers (based on 3868 papers).***

| Landscape elements categories | Included landscapes elements' description |
|---|---|
| Diverse floral resources | floral stripes and patches within orchards, rich diverse native and wildflower floral strips, flowering ground cover, diverse floral field margins, native and non native flowering herbaceous strips adjacent fields. |
| Complex diverse SNH | mixed of different elements: small patches of forest , pasture, grassland, shrubland surrounding, complex hedgerows with grassy elements or woody and rich floral understory species, prairie strips, diverse rich field margins with higher native and introduced plants and trees, diverse field border species, agri-environment schemes(e.g. wildflower strips/areas, grassy field margins, organic farming), vegetation cover in inter-row, mixed plantation, heterogeneous diverse SNH, forest edges, green spaces and domestic gardens, zoos, diverse urban parks, public parks, castle parks, disaggregated forested area, small water bodies, open lawns and trees, zoned woody, shrubby and grassy buffers, wetland, diverse ground cover. |
| Complex diverse NH | Diverse natural species, heterogenous natural buffer, native forested, shrubs, grassy buffer, natural forest, diverse native natural habitat fragments in mosaic landscapes, old natural field margins, natural heterogeneous vegetation cover, vegetation cover with diverse native deep rooted species with more reinforcing effect and low surcharge, natural young trees |
| Woody elements | Silvoarable alleys, agroforestry, narrow woodland corridors,woody corridors that fragment fields, woody vegetation margins, hedgerows, trees canopy cover, street trees, native shrubland, hardwood buffer, native diverse woody buffer. |
| Grassy elements | patches of grassland surrounding, diverse SN pasture and field border, diverse rich patches of urban meadow |
| Forest | forest cover surrounding, forest edges associated with fallow or hedges, forest patches surrounding, forest corridors, diverse forested native and non native buffer, deep rooted natural forest species |

***Table 2. Land cover classification***

| **Land cover classification** | **Binary classification** |
|---|---|
| tree cover | 1 |
| shrubland | 1 |
| natural grassland | 1 |
| herbaceous wetland | 1 |
| moss and lichen | 1 |
| mangroves | 1 |
| cropland | 0 |
| built-up areas | 0 |
| pastureland | 0 |
| bare/sparse vegetation | NA |
| snow and ice | NA |
| permanent water bodies | NA |

***Table 3. Spatial distribution of the functional integrity thresholds.*** *Integrity is calculated as the average value of the binary classified layer (natural/human-modified) within a 1 km$^2$ radius for (1) human-modified lands and (2) the total global land surface. We performed an additional sensitivity analysis for the human-modified lands calculation in which we use additional data to more explicitly account for plantations as part of the human-modified landscape. For full methodological details see the Supplementary Information.*

| **Integrity threshold applied (%)** | **Percent land above integrity threshold** | | |
|---|---|---|---|
| | **Functionally intact human-modified lands (%)** | **Functionally intact human-modified lands sensitivity analysis (%)** | **Functionally intact global land surface (%)** |
| ***10*** | ***49*** | ***53*** | ***63*** |
| ***15*** | ***41*** | ***45*** | ***62*** |
| ***20*** | ***36*** | ***39*** | ***62*** |
| ***25*** | ***31*** | ***34*** | ***61*** |
| 30 | 27 | 29 | 61 |

| | | | |
|---|---|---|---|
| 40 | 20 | 21 | 60 |
| 50 | 14 | 15 | 58 |
| 60 | 10 | 10 | 57 |
| 70 | 6 | 6 | 56 |
| 80 | 4 | 4 | 55 |
| 90 | 2 | 2 | 53 |
| 100 | 0 | 0 | 47 |

### 6. References used in the NCP analysis

1. Adhikari, S., Adhikari, A., Weaver, D. K., Bekkerman, A. & Menalled, F. D. Impacts of Agricultural Management Systems on Biodiversity and Ecosystem Services in Highly Simplified Dryland Landscapes. SUSTAINABILITY 11, (2019).
2. Aguiar Jr., T. R., Rasera, K., Parron, L. M., Brito, A. G. & Ferreira, M. T. Nutrient removal effectiveness by riparian buffer zones in rural temperate watersheds: The impact of no-till crops practices. Agric. Water Manag. 149, 74–80 (2015).
3. Aguilera, G. et al. Crop diversity benefits carabid and pollinator communities in landscapes with semi-natural habitats. J. Appl. Ecol. 57, 2170–2179 (2020).
4. Albrecht, M. et al. The effectiveness of flower strips and hedgerows on pest control, pollination services and crop yield: a quantitative synthesis. Ecol. Lett. 23, 1488–1498 (2020).

5. Alignier, A. & Aviron, S. Time-lagged response of carabid species richness and composition to past management practices and landscape context of semi-natural field margins. J. Environ. Manage. 204, 282–290 (2017).
6. Aristizábal, N. & Metzger, J. P. Landscape structure regulates pest control provided by ants in sun coffee farms. J. Appl. Ecol. 56, 21–30 (2019).
7. Arnold, S. E. J. et al. Beneficial insects are associated with botanically rich margins with trees on small farms. Sci. Rep. 11, 15190 (2021).
8. Arroyo-Rodríguez, V. et al. Designing optimal human-modified landscapes for forest biodiversity conservation. Ecol. Lett. 23, 1404–1420 (2020).
9. Avelino, J., Romero-Gurdián, A., Cruz-Cuellar, H. F. & Declerck, F. A. Landscape context and scale differentially impact coffee leaf rust, coffee berry borer, and coffee root-knot nematodes. Ecol. Appl. 22, 584–596 (2012).
10. Aviron, S., Lalechere, E., Duflot, R., Parisey, N. & Poggi, S. Connectivity of cropped vs. semi-natural habitats mediates biodiversity: A case study of carabid beetles communities. Agric. Ecosyst. Environ. 268, 34–43 (2018).
11. Badenhausser, I. et al. Increasing amount and quality of green infrastructures at different scales promotes biological control in agricultural landscapes. Agric. Ecosyst. Environ. 290, 106735 (2020).
12. Bailey, S. et al. Distance from forest edge affects bee pollinators in oilseed rape fields. Ecol. Evol. 4, 370–380 (2014).
13. Barboza, E. P. et al. Green space and mortality in European cities: a health impact assessment study. Lancet Planet. Health 5, e718–e730 (2021).
14. Bartual, A. M. et al. The potential of different semi-natural habitats to sustain pollinators and natural enemies in European agricultural landscapes. Agric. Ecosyst. Environ. 279, 43–52 (2019).
15. Beegum, S., Jainet, P., Emil, D., Sudheer, K. & Das, S. Integrated Simulation Modeling Approach for Investigating Pore Water Pressure Induced Landslides. (2022).
16. Bergman, K.-O., Dániel-Ferreira, J., Milberg, P., Öckinger, E. & Westerberg, L. Butterflies in Swedish grasslands benefit from forest and respond to landscape composition at different spatial scales. Landsc. Ecol. 33, 2189–2204 (2018).
17. Cao, X., Song, C., Xiao, J. & Zhou, Y. The Optimal Width and Mechanism of Riparian Buffers for Storm Water Nutrient Removal in the Chinese Eutrophic Lake Chaohu Watershed. Water 10, (2018).
18. Carvalheiro, L. G., Seymour, C. L., Veldtman, R. & Nicolson, S. W. Pollination services decline with distance from natural habitat even in biodiversity-rich areas. J. Appl. Ecol. 47, 810–820 (2010).
19. Chang, K. G., Sullivan, W. C., Lin, Y.-H., Su, W. & Chang, C.-Y. The effect of biodiversity on green space users' wellbeing—An empirical investigation using physiological evidence. Sustainability 8, 1049 (2016).
20. Chaplin-Kramer, R., O'Rourke, M. E., Blitzer, E. J. & Kremen, C. A meta-analysis of crop pest and natural enemy response to landscape complexity. Ecol. Lett. 14 9, 922–32 (2011).
21. Chen, J. et al. Threshold effects of vegetation coverage on soil erosion control in small watersheds of the red soil hilly region in China. Ecol. Eng. 132, 109–114 (2019).
22. Christen, B. & Dalgaard, T. Buffers for biomass production in temperate European agriculture: A review and synthesis on function, ecosystem services and implementation. Biomass Bioenergy 55, 53–67 (2013).

23. Clough, Y., Kirchweger, S. & Kantelhardt, J. Field sizes and the future of farmland biodiversity in European landscapes. Conserv. Lett. 13, e12752 (2020).
24. Cole, L. J., Stockan, J. & Helliwell, R. Managing riparian buffer strips to optimise ecosystem services: A review. Agric. Ecosyst. Environ. 296, 106891 (2020).
25. Cox, D. T. C. et al. Doses of Nearby Nature Simultaneously Associated with Multiple Health Benefits. Int. J. Environ. Res. Public. Health 14, (2017).
26. Dainese, M., Luna, D. I., Sitzia, T. & Marini, L. Testing scale-dependent effects of seminatural habitats on farmland biodiversity. Ecol. Appl. 25, 1681–1690 (2015).
27. Dala-Corte, R. B. et al. Thresholds of freshwater biodiversity in response to riparian vegetation loss in the Neotropical region. J. Appl. Ecol. 57, 1391–1402 (2020).
28. Dassou, A. G. & Tixier, P. Response of pest control by generalist predators to local-scale plant diversity: a meta-analysis. Ecol. Evol. 6, 1143–1153 (2016).
29. Deeks, L., Duzant, J., Owens, P. & Wood, G. A decision support framework for effective design and placement of vegetated buffer strips within agricultural field systems. Adv. Agron. 114, 225–248 (2012).
30. Douglas, G. et al. Reducing shallow landslide occurrence in pastoral hill country using wide-spaced trees. Land Degrad. Dev. 24, 103–114 (2013).
31. Douglas-Mankin, K. R., Helmers, M. J. & Harmel, R. D. Review of Filter Strip Performance and Function for Improving Water Quality from Agricultural Lands. Trans. ASABE 64, 659–674 (2021).
32. Duarte, G. T., Santos, P. M., Cornelissen, T. G., Ribeiro, M. C. & Paglia, A. P. The effects of landscape patterns on ecosystem services: meta-analyses of landscape services. Landsc. Ecol. 33, 1247–1257 (2018).
33. Emadi-Tafti, M., Ataie-Ashtiani, B. & Hosseini, S. M. Integrated impacts of vegetation and soil type on slope stability: A case study of Kheyrud Forest, Iran. Ecol. Model. 446, (2021).
34. Fountain, M. T. Impacts of Wildflower Interventions on Beneficial Insects in Fruit Crops: A Review. Insects 13, (2022).
35. Gagic, V., Paull, C. & Schellhorn, N. A. Ecosystem service of biological pest control in Australia: the role of non-crop habitats within landscapes. AUSTRAL Entomol. 57, 194–206 (2018).
36. Garibaldi, L. A. et al. Working landscapes need at least 20% native habitat. Conserv. Lett. 14, e12773 (2021).
37. Garibaldi, L. et al. Stability of pollination services decreases with isolation from natural areas despite honey bee visits. Ecol. Lett. 14, 1062–72 (2011).
38. Garratt, M. P. D., Senapathi, D., Coston, D. J., Mortimer, S. R. & Potts, S. G. The benefits of hedgerows for pollinators and natural enemies depends on hedge quality and landscape context. Agric. Ecosyst. Environ. 247, 363–370 (2017).
39. Genet, M. et al. Root reinforcement in plantations of Cryptomeria japonica D. Don: effect of tree age and stand structure on slope stability. For. Ecol. Manag. 256, 1517–1526 (2008).
40. Genet, M., Stokes, A., Fourcaud, T. & Norris, J. E. The influence of plant diversity on slope stability in a moist evergreen deciduous forest. Spec. Issue Veg. Slope Stab. 36, 265–275 (2010).
41. Geslin, B. et al. Spatiotemporal changes in flying insect abundance and their functional diversity as a function of distance to natural habitats in a mass flowering crop. Agric. Ecosyst. Environ. 229, 21–29 (2016).

42. Guidotti, V. et al. Changes in Brazil's Forest Code can erode the potential of riparian buffers to supply watershed services. Land Use Policy 94, 104511 (2020).
43. Gyssels, G., Poesen, J., Bochet, E. & Li, Y. Impact of plant roots on the resistance of soils to erosion by water: a review. Prog. Phys. Geogr. 29, 189–217 (2005).
44. Ha, J., Kim, H. J. & With, K. A. Urban green space alone is not enough: A landscape analysis linking the spatial distribution of urban green space to mental health in the city of Chicago. Landsc. Urban Plan. 218, 104309 (2022).
45. Haan, N. L., Zhang, Y. & Landis, D. A. Predicting Landscape Configuration Effects on Agricultural Pest Suppression. TRENDS Ecol. Evol. 35, 175–186 (2020).
46. Hansen, B. D., Reich, P., Cavagnaro, T. R. & Lake, P. S. Challenges in applying scientific evidence to width recommendations for riparian management in agricultural Australia. Ecol. Manag. Restor. 16, 50–57 (2015).
47. Hansen, B., Reich, P. & Cavagnaro, T. Minimum width requirements for riparian zones to protect flowing waters and to conserve biodiversity: a review and recommendations With application to the State of Victoria. (2010).
48. Hass, A. L. et al. Landscape configurational heterogeneity by small-scale agriculture, not crop diversity, maintains pollinators and plant reproduction in western Europe. Proc. R. Soc. B Biol. Sci. 285, 20172242 (2018).
49. Hill, A. R. Groundwater nitrate removal in riparian buffer zones: a review of research progress in the past 20 years. Biogeochemistry 143, 347–369 (2019).
50. Hill, A. R., Devito, K. J. & Vidon, P. G. Long-term nitrate removal in a stream riparian zone. Biogeochemistry 121, 425–439 (2014).
51. Holland, J. M. et al. Structure, function and management of semi-natural habitats for conservation biological control: a review of European studies. Pest Manag. Sci. 72, 1638–1651 (2016).
52. Holz, D. J., Williard, K. W., Edwards, P. J. & Schoonover, J. E. Soil erosion in humid regions: a review. J. Contemp. Water Res. Educ. 154, 48–59 (2015).
53. Hou, G. et al. A vegetation configuration pattern with a high-efficiency purification ability for TN, TP, AN, AP, and COD based on comprehensive assessment results. Sci. Rep. 9, 2427 (2019).
54. Iñiguez–Armijos, C., Leiva, A., Frede, H., Hampel, H. & Breuer, L. Deforestation and Benthic Indicators: How Much Vegetation Cover Is Needed to Sustain Healthy Andean Streams? PLOS ONE 9, e105869 (2014).
55. Jiang, B., Chang, C.-Y. & Sullivan, W. C. A dose of nature: Tree cover, stress reduction, and gender differences. Landsc. Urban Plan. 132, 26–36 (2014).
56. Jiang, F., Preisendanz, H. E., Veith, T. L., Cibin, R. & Drohan, P. J. Riparian buffer effectiveness as a function of buffer design and input loads. J. Environ. Qual. 49, 1599–1611 (2020).
57. Johnson, S. R., Burchell, M. R., Evans, R. O., Osmond, D. L. & Gilliam, J. W. Riparian buffer located in an upland landscape position does not enhance nitrate-nitrogen removal. Ecol. Eng. 52, 252–261 (2013).
58. Joseph, J. et al. A spatially extended model to assess the role of landscape structure on the pollination service of Apis mellifera. Ecol. Model. 431, 109201 (2020).
59. Karp Daniel S. et al. Crop pests and predators exhibit inconsistent responses to surrounding landscape composition. Proc. Natl. Acad. Sci. 115, E7863–E7870 (2018).
60. Kennedy, C. M. et al. A global quantitative synthesis of local and landscape effects on wild bee pollinators in agroecosystems. Ecol. Lett. 16, 584–599 (2013).

61. Klaus, F., Tscharntke, T., Uhler, J. & Grass, I. Calcareous grassland fragments as sources of bee pollinators for the surrounding agricultural landscape. Glob. Ecol. Conserv. 26, e01474 (2021).
62. Klemas, V. Remote sensing of riparian and wetland buffers: an overview. J. Coast. Res. 30, 869–880 (2014).
63. Kline, O. & Joshi, N. K. Mitigating the Effects of Habitat Loss on Solitary Bees in Agricultural Ecosystems. Agriculture 10, (2020).
64. Kolb, S., Uzman, D., Leyer, I., Reineke, A. & Entling, M. H. Differential effects of semi-natural habitats and organic management on spiders in viticultural landscapes. Agric. Ecosyst. Environ. 287, 106695 (2020).
65. Kondo, M. C. et al. Health impact assessment of Philadelphia's 2025 tree canopy cover goals. Lancet Planet. Health 4, e149–e157 (2020).
66. Konijnendijk, C. The 3-30-300 Rule for Urban Forestry and Greener Cities. IUCN Urban Alliance.
67. Kormann, U. et al. Local and landscape management drive trait-mediated biodiversity of nine taxa on small grassland fragments. Divers. Distrib. 21, 1204–1217 (2015).
68. Kral-O'Brien, K. C., O'Brien, P. L., Hovick, T. J. & Harmon, J. P. Meta-analysis: Higher Plant Richness Supports Higher Pollinator Richness Across Many Land Use Types. Ann. Entomol. Soc. Am. 114, 267–275 (2021).
69. Kremen, C. Ecological intensification and diversification approaches to maintain biodiversity, ecosystem services and food production in a changing world. Emerg. Top. LIFE Sci. 4, 229–240 (2020).
70. Labrière, N., Locatelli, B., Laumonier, Y., Freycon, V. & Bernoux, M. Soil erosion in the humid tropics: A systematic quantitative review. Agric. Ecosyst. Environ. 203, 127–139 (2015).
71. Lajos, K., Samu, F., Bihaly, Á. D., Fülöp, D. & Sárospataki, M. Landscape structure affects the sunflower visiting frequency of insect pollinators. Sci. Rep. 11, 8147 (2021).
72. Larsen, A. E. & Noack, F. Impact of local and landscape complexity on the stability of field-level pest control. Nat. Sustain. 4, 120–128 (2021).
73. Larson, L. R., Jennings, V. & Cloutier, S. A. Public parks and wellbeing in urban areas of the United States. PLoS One 11, e0153211 (2016).
74. Lee, P., Smyth, C. & Boutin, S. Quantitative review of riparian buffer width guidelines from Canada and the United States. J. Environ. Manage. 70, 165–180 (2004).
75. Letourneau, D. et al. Does plant diversity benefit agroecosystems? A synthetic review. Ecol. Appl. Publ. Ecol. Soc. Am. 21, 9–21 (2011).
76. Liivamägi, A., Kuusemets, V., Kaart, T., Luig, J. & Diaz-Forero, I. Influence of habitat and landscape on butterfly diversity of semi-natural meadows within forest-dominated landscapes. J. Insect Conserv. 18, 1137–1145 (2014).
77. Lin, X., Tang, J., Li, Z. & Li, H. Finite Element Simulation of Total Nitrogen Transport in Riparian Buffer in an Agricultural Watershed. Sustainability 8, (2016).
78. Lind, L., Hasselquist, E. M. & Laudon, H. Towards ecologically functional riparian zones: A meta-analysis to develop guidelines for protecting ecosystem functions and biodiversity in agricultural landscapes. J. Environ. Manage. 249, 109391 (2019).
79. Liu, J. et al. The effects of vegetation on runoff and soil loss: Multidimensional structure analysis and scale characteristics. J. Geogr. Sci. 28, 59–78 (2018).
80. Luke, S. H. et al. Riparian buffers in tropical agriculture: Scientific support, effectiveness and directions for policy. J. Appl. Ecol. 56, 85–92 (2019).

81. Lv, J. & Wu, Y. Nitrogen removal by different riparian vegetation buffer strips with different stand densities and widths. Water Supply 21, 3541–3556 (2021).
82. Lyu, C. et al. Nitrogen retention effect of riparian zones in agricultural areas: A meta-analysis. J. Clean. Prod. 315, 128143 (2021).
83. Mancuso, G., Bencresciuto, G. F., Lavrnić, S. & Toscano, A. Diffuse Water Pollution from Agriculture: A Review of Nature-Based Solutions for Nitrogen Removal and Recovery. Water 13, (2021).
84. Mankin, K. R., Ngandu, D. M., Barden, C. J., Hutchinson, S. L. & Geyer, W. A. Grass-Shrub Riparian Buffer Removal of Sediment, Phosphorus, and Nitrogen From Simulated Runoff1. JAWRA J. Am. Water Resour. Assoc. 43, 1108–1116 (2007).
85. Marja, R., Tscharntke, T. & Batáry, P. Increasing landscape complexity enhances species richness of farmland arthropods, agri-environment schemes also abundance – A meta-analysis. Agric. Ecosyst. Environ. 326, 107822 (2022).
86. Marselle, M. R. et al. Urban street tree biodiversity and antidepressant prescriptions. Sci. Rep. 10, 22445 (2020).
87. Martin, E., Seo, B., Park, C., Reineking, B. & Steffan-Dewenter, I. Scale-dependent effects of landscape composition and configuration on natural enemy diversity, crop herbivory, and yields. Ecol. Appl. 26, 448–462 (2016).
88. Martin, E. A. et al. The interplay of landscape composition and configuration: new pathways to manage functional biodiversity and agroecosystem services across Europe. Ecol. Lett. 22, 1083–1094 (2019).
89. Mayer, P. M., Reynolds Jr, S. K., McCutchen, M. D. & Canfield, T. J. Meta-analysis of nitrogen removal in riparian buffers. J. Environ. Qual. 36, 1172–1180 (2007).
90. Mazzei, M. P., Vesprini, J. L. & Galetto, L. Seminatural habitats and their proximity to the crop enhances canola (Brassica napus) pollination and reproductive parameters in Argentina. Crop Sci. 61, 2713–2721 (2021).
91. Moreira, T. C. L. et al. Assessing the impact of urban environment and green infrastructure on mental health: results from the São Paulo Megacity Mental Health Survey. J. Expo. Sci. Environ. Epidemiol. 32, 205–212 (2022).
92. Muchane, M. N. et al. Agroforestry boosts soil health in the humid and sub-humid tropics: A meta-analysis. Agric. Ecosyst. Environ. 295, 106899 (2020).
93. Mueller, N. et al. Changing the urban design of cities for health: The superblock model. Environ. Int. 134, 105132 (2020).
94. Nagy, R. K., Bell, L. W., Schellhorn, N. A. & Zalucki, M. P. Role of grasslands in pest suppressive landscapes: how green are my pastures? Austral Entomol. 59, 227–237 (2020).
95. Olafsdottir, G. et al. Health Benefits of Walking in Nature: A Randomized Controlled Study Under Conditions of Real-Life Stress. Environ. Behav. 52, 248–274 (2020).
96. Oldén, A., Selonen, V. A. O., Lehkonen, E. & Kotiaho, J. S. The effect of buffer strip width and selective logging on streamside plant communities. BMC Ecol. 19, 9 (2019).
97. Otieno, M. et al. Enhancing legume crop pollination and natural pest regulation for improved food security in changing African landscapes. Glob. Food Secur. 26, 100394 (2020).
98. Pardo, A. & Borges, P. A. V. Worldwide importance of insect pollination in apple orchards: A review. Agric. Ecosyst. Environ. 293, 106839 (2020).

99. Perrot, T., Rusch, A., Coux, C., Gaba, S. & Bretagnolle, V. Proportion of Grassland at Landscape Scale Drives Natural Pest Control Services in Agricultural Landscapes. Front. Ecol. Evol. 9, (2021).
100. Phogat, V. et al. Optimizing the riparian zone width near a river for controlling lateral migration of irrigation water and solutes. J. Hydrol. 570, 637–646 (2019).
101. Plećaš, M. et al. Landscape composition and configuration influence cereal aphid–parasitoid–hyperparasitoid interactions and biological control differentially across years. Agric. Ecosyst. Environ. 183, 1–10 (2014).
102. Prasetyo, A., Setyawan, C. & Tirtalistyani, R. Vegetation cover modelling for soil erosion control in agricultural watershed. in vol. 653 012033 (IOP Publishing, 2021).
103. Prosser, R. S. et al. A review of the effectiveness of vegetated buffers to mitigate pesticide and nutrient transport into surface waters from agricultural areas. J. Environ. Manage. 261, 110210 (2020).
104. Quinton, J. N., Edwards, G. & Morgan, R. The influence of vegetation species and plant properties on runoff and soil erosion: results from a rainfall simulation study in south east Spain. Soil Use Manag. 13, 143–148 (1997).
105. Rahimi, E., Barghjelveh, S. & Dong, P. Estimating landscape structure effects on pollination for management of agricultural landscapes. Ecol. Process. 10, 59 (2021).
106. Rahman, M. A. et al. Spatial and temporal changes of outdoor thermal stress: influence of urban land cover types. Sci. Rep. 12, 671 (2022).
107. Ramesh, R., Kalin, L., Hantush, M. & Chaudhary, A. A secondary assessment of sediment trapping effectiveness by vegetated buffers. Ecol. Eng. 159, 106094 (2021).
108. Ratto, F. et al. Proximity to natural habitat and flower plantings increases insect populations and pollination services in South African apple orchards. J. Appl. Ecol. 58, 2540–2551 (2021).
109. Redlich, S., Martin, E. A. & Steffan-Dewenter, I. Landscape-level crop diversity benefits biological pest control. J. Appl. Ecol. 55, 2419–2428 (2018).
110. Rossi, L. M. W. et al. Sensitivity of the landslide model LAPSUS_LS to vegetation and soil parameters. Soil Bio- Eco-Eng. Use Veg. Improve Slope Stab. - Proc. Fourth Int. Conf. 109, 249–255 (2017).
111. Rusch, A., Bommarco, R., Jonsson, M., Smith, H. G. & Ekbom, B. Flow and stability of natural pest control services depend on complexity and crop rotation at the landscape scale. J. Appl. Ecol. 50, 345–354 (2013).
112. Rusch, A. et al. Agricultural landscape simplification reduces natural pest control: A quantitative synthesis. Agric. Ecosyst. Environ. 221, 198–204 (2016).
113. Saturni, F. T., Jaffé, R. & Metzger, J. P. Landscape structure influences bee community and coffee pollination at different spatial scales. Agric. Ecosyst. Environ. 235, 1–12 (2016).
114. Scheper, J. et al. Environmental factors driving the effectiveness of European agri-environmental measures in mitigating pollinator loss – a meta-analysis. Ecol. Lett. 16, 912–920 (2013).
115. Shackelford, G. et al. Comparison of pollinators and natural enemies: a meta-analysis of landscape and local effects on abundance and richness in crops. Biol. Rev. 88, 1002–1021 (2013).
116. Shanahan, D. F., Lin, B. B., Gaston, K. J., Bush, R. & Fuller, R. A. What is the role of trees and remnant vegetation in attracting people to urban parks? Landsc. Ecol. 30, 153–165 (2015).

135. Wang, Z.-J., Jiao, J.-Y., Rayburg, S., Wang, Q.-L. & Su, Y. Soil erosion resistance of "Grain for Green" vegetation types under extreme rainfall conditions on the Loess Plateau, China. CATENA 141, 109–116 (2016).
136. Watson, J. C., Wolf, A. T. & Ascher, J. S. Forested Landscapes Promote Richness and Abundance of Native Bees (Hymenoptera: Apoidea: Anthophila) in Wisconsin Apple Orchards. Environ. Entomol. 40, 621–632 (2011).
137. Wenzel, A., Grass, I., Belavadi, V. V. & Tscharntke, T. How urbanization is driving pollinator diversity and pollination – A systematic review. Biol. Conserv. 241, 108321 (2020).
138. White, M. P. et al. Spending at least 120 minutes a week in nature is associated with good health and wellbeing. Sci. Rep. 9, 7730 (2019).
139. Winter, S. et al. Effects of vegetation management intensity on biodiversity and ecosystem services in vineyards: A meta-analysis. J. Appl. Ecol. 55, 2484–2495 (2018).
140. Wood, E. et al. Not all green space is created equal: Biodiversity predicts psychological restorative benefits from urban green space. Front. Psychol. 9, 2320 (2018).
141. Woodcock, B. A. et al. Spill-over of pest control and pollination services into arable crops. Agric. Ecosyst. Environ. 231, 15–23 (2016).
142. World Health Organization. Urban green spaces and health. (2016).
143. Wu, G.-L. et al. Trade-off between vegetation type, soil erosion control and surface water in global semi-arid regions: A meta-analysis. J. Appl. Ecol. 57, 875–885 (2020).
144. Wu, P. et al. Improving Habitat Quality at the Local and Landscape Scales Increases Wild Bee Assemblages and Associated Pollination Services in Apple Orchards in China. Front. Ecol. Evol. 9, (2021).
145. Xu, C. et al. Runoff and soil erosion responses to rainfall and vegetation cover under various afforestation management regimes in subtropical montane forest. Land Degrad. Dev. 30, 1711–1724 (2019).
146. Yan, R., Zhang, X., Yan, S. & Chen, H. Estimating soil erosion response to land use/cover change in a catchment of the Loess Plateau, China. Int. Soil Water Conserv. Res. 6, 13–22 (2018).
147. Zamorano, J., Bartomeus, I., Grez, A. A. & Garibaldi, L. A. Field margin floral enhancements increase pollinator diversity at the field edge but show no consistent spillover into the crop field: a meta-analysis. Insect Conserv. Divers. 13, 519–531 (2020).
148. Zhang, X., Song, J., Wang, Y., Sun, H. & Li, Q. Threshold effects of vegetation coverage on runoff and soil loss in the Loess Plateau of China: A meta-analysis. Geoderma 412, 115720 (2022).
149. Zhang, X., Liu, X., Zhang, M., Dahlgren, R. A. & Eitzel, M. A Review of Vegetated Buffers and a Meta-analysis of Their Mitigation Efficacy in Reducing Nonpoint Source Pollution. J. Environ. Qual. 39, 76–84 (2010).
150. Zheng, Y., Wang, H., Qin, Q. & Wang, Y. Effect of plant hedgerows on agricultural non-point source pollution: a meta-analysis. Environ. Sci. Pollut. Res. 27, 24831–24847 (2020).
151. Zhongming, W., Lees, B. G., Feng, J., Wanning, L. & Haijing, S. Stratified vegetation cover index: A new way to assess vegetation impact on soil erosion. CATENA 83, 87–93 (2010).
152. Zuazo, V. H. D. & Pleguezuelo, C. R. R. Soil-erosion and runoff prevention by plant covers: a review. Sustain. Agric. 785–811 (2009).

153. Zurbuchen, A. et al. Maximum foraging ranges in solitary bees: only few individuals have the capability to cover long foraging distances. Biol. Conse*rv. 143, 669–676 (2010).*
